# High ammonium suppresses nitrate transporter expression and constrains thermal plasticity in the green macroalga *Codium cylindricum*

**DOI:** 10.64898/2026.09.10.750591

**Authors:** Kai-Le Zhong, Pamela A. Fernandez, Bayden D. Russell, Juan Diego Gaitan-Espitia

## Abstract

Nitrogen source and availability can modulate thermal stress responses in marine macroalgae, yet the transcriptional mechanisms underpinning these effects in green seaweeds remain poorly understood. We tested how three N sources (ammonium, nitrate, urea) at low (5 μM) and high (100 μM) concentrations influence the thermal physiology and gene expression of *Codium cylindricum*. Thermal performance curves were constructed for growth, photosynthesis, and pigment content across a 5–29 °C gradient, coupled with transcriptomic analysis at three key temperatures. At high concentrations, ammonium reduced maximum growth rate by 25% and lowered the thermal optimum relative to nitrate and urea, while nitrate supported the broadest thermal performance. Urea constrained thermal breadth regardless of concentration. These responses were linked to differential expression of Nitrogen assimilation genes. Ammonium suppressed nitrate transporter genes (*NRT*) and nitrite transporter (*FNT*), and drove pigment accumulation with rising temperature despite reduced growth, suggesting resource reallocation from growth to photoprotection. In contrast, nitrate upregulated photosystem and phosphorylation-related genes. Our findings demonstrate that Nitrogen source and concentration jointly determine the thermal threshold of *C. cylindricum*, with high ammonium reducing both maximum growth and the temperature at which it is achieved. This highlights how shifting coastal nutrient regimes may alter macroalgal resilience under ocean warming.

**Highlight:** In the green macroalga *Codium cylindricum*, high ammonium availability suppresses growth and constrains thermal plasticity through downregulation of nitrate assimilation genes, whereas nitrate supports broader thermal performance. Urea limits thermal breadth regardless of concentration.

## 1 Introduction

Macroalgae are ecologically and economically important marine primary producers that play central roles in the functional and structural dynamics of coastal ecosystems worldwide (Hurd et al. 2014). These organisms experience dynamic environments, where temperature has been a critical factor influencing both their ecology and evolution (Eggert 2012; Raven et al. 2002). Seasonal and temporal changes in temperature modulate intra-and inter-specific patterns in the physiology (Wernberg et al. 2016), phenology (Wiencke et al. 2009), and geographic distribution (Bartsch et al. 2012; Zhong et al. 2024) of these primary producers. Such patterns have often been linked to local adaptation and organismal physiological and molecular plasticity (i.e., phenotypic plasticity) (Flukes et al. 2015; Machado-Monteiro et al. 2019; Bennett et al. 2021; Liu et al. 2022). A central framework for quantifying this plasticity is the thermal performance curve (TPC), which predicts that fitness (or other performance traits) increases with rising temperature up to a maximum (r_max) at an optimal temperature (T_opt), beyond which further warming causes inhibition and declines in performance (Schulte et al. 2011; Huey & Stevenson, 1979). The shape of the TPC is determined by parameters including r_max, T_opt, and thermal breadth (T_br), which collectively capture both vertical shifts in maximum performance and horizontal shifts in thermal range (Fernandez et al. 2020).

However, plastic capacities to tolerate thermal variation and stress can be influenced by co-occurring environmental factors such as salinity (Monteiro et al. 2021), irradiance (Paine et al. 2021), and nutrient availability (Huo et al. 2021; Fernández et al. 2020). Therefore, to better understand ecological dynamics and potential evolutionary responses of macroalgae to temperature changes (e.g., seasonality, ocean warming, heatwaves), it is important to assess the influence of co-occurring factors on thermal tolerances and the underlying mechanistic regulation.

As with other primary producers, nitrogen (N) is an essential nutrient in macroalgae, supporting developmental (e.g., growth, reproduction) and functional (e.g., enzymatic activity, pigment formation, photosynthesis) processes (Chen et al. 2023; Fernández et al. 2020). In coastal-marine systems, N is available in three primary forms: ammonium (NH□□), nitrate (NO□□), and urea (CO(NH□)□) (Abreu et al. 2011). The two inorganic forms (i.e., NH and NO□□) are considered the main N sources used by macroalgae, with most species showing higher affinity for NH due to the lower energetic costs associated with its direct uptake and assimilation (Hurd et al. 2014; Smith et al. 2019; Fan et al. 2020; Zhang et al. 2026). However, the affinity, uptake rate, utilization, and assimilation of these N forms can vary across macroalgal species, depending on factors such as size (Hein et al. 1995), physiological status (Roleda & Hurd, 2019), and local environmental availability (Harrison & Hurd, 2001; Ale et al. 2011; Han et al. 2018; Smith et al. 2019). For example, under varying ammonium/nitrate ratios, *Caulerpa lentillifera* selectively takes up nitrate at high rates (7.43–50.43 μmol g□¹ dw h□¹), whereas *Gracilaria lichenoides* preferentially takes up ammonium (10.27–14.14 μmol g ¹ dw h ¹), with assimilation rates for both forms shifting significantly with nutrient availability in both species (Liu et al. 2016).

Changes in N source availability can trigger contrasting physiological responses in macroalgae, reflecting species-specific differences in nutritional requirements, N source preferences, and tolerances to high or low N concentrations (Rabalais et al. 2009; Sarmiento et al. 1993; Herbert, 1999; Pedersen et al. 1997; Korb & Gerard, 2000). For example, in some species (i.e., *Sargassum horneri*), high NH□□ concentrations inhibit growth and survival (Miki et al. 2016), while in others (e.g., *Ulva australis*), NH□□ enrichment increases productivity and growth (Reidenbach et al. 2017). N availability can also modulate thermal responses in macroalgae. In some brown algae (e.g., *Macrocystis pyrifera*, *Fucus vesiculosus*), N enrichment in the form of NO□□ has been shown to enhance thermal tolerance (Fernández et al. 2020) and buffer thermal stress (Schmid et al. 2020; Colvard and Helmuth, 2017). While these findings underscore the significance of N enrichment in modulating thermal responses, most of this research has been conducted in economically important brown macroalgae. Consequently, our understanding of how green macroalgae, especially beyond the genus *Ulva*, respond to N enrichment in relation to thermal stress remains limited. Furthermore, the regulatory mechanisms underlying these responses, as well as their capacity for plastic adjustments in relation to different N forms and availability, remain poorly understood in green macroalgae. Gaining insight into these mechanisms and their role in modulating thermal physiological responses is essential for improving our understanding of the adaptive capacity and ecological resilience of macroalgae to ocean warming and extreme temperature events.

Blooms and invasive events of *Codium* species have been frequently recorded in nutrient-enriched subtropical and tropical coastal habitats (Lapointe, 1997, 2005; Israel et al. 2010; Chang et al. 2010). Sufficient nitrogen supply supports the fast-growing strategy of these macroalgae, suggesting that nitrogen availability and form could profoundly shape their thermal physiological characteristics and overall ecological competitiveness. However, the mechanistic link between N regime and thermal physiology in Codium species remains unexplored. Here, we assessed how N sources (ammonium, nitrate, and urea) and their availability influence the thermal physiology (thermal tolerances and plasticity) and associated transcriptional regulation of the green macroalga *Codium cylindricum*. We employed the thermal performance curve (TPC) framework, which quantifies how performance traits vary with temperature and captures key parameters including the maximum performance (r_max), optimal temperature (T_opt), thermal breadth (T_br), and critical thermal limits (CT_min and CT_max) (Schulte et al. 2011; Huey & Stevenson, 1979). Shifts in these parameters reflect changes in thermal plasticity, with vertical shifts indicating altered maximum performance and horizontal shifts indicating changes in the thermal range (Fernández et al. 2020). We constructed TPCs for growth, photosynthesis, and pigment content across a 5–29 °C gradient and coupled these with transcriptomic analysis at three key temperatures to identify molecular pathways underpinning N-source-dependent thermal responses. We hypothesized that distinct N sources and concentrations would alter the thermal physiology of *C. cylindricum*. Specifically, we predicted that high N concentrations would enhance thermal tolerance and elevate optimal temperature, regardless of N source. We further predicted that these differential responses would be regulated by changes in the expression of key genes involved in metabolic pathways (e.g., N metabolism and fatty acid biosynthesis), oxidative stress, photorespiration, and photosynthesis (Liu et al. 2019; Li et al. 2019). Our study provides insights into N speciation and how it modulates thermal resilience in macroalgae, advancing our understanding of their adaptive capacity under the concurrent pressures of ocean warming and coastal eutrophication.

## 2 Materials and methods

### 2.1 Sample collection and species identification

Thalli of *Codium cylindricum* were collected from Little Palm Beach (22.32°N, 114.28°E), Hong Kong, China. Samples were immediately transported to the laboratory at the University of Hong Kong in dark insulated containers filled with ambient seawater. Samples were gently cleaned with filtered seawater (0.22μm) to remove the visible fouling and epibionts. Each sample of about 5 g of fresh weight (FW) was acclimated under controlled laboratory conditions (17 ± 1 °C, 35ppt, 50∼60 μmol photons m^−2^ s^−1^ (PAR), 12:12 h light: dark period) with UV-filtered seawater and modified Von Stosch Enriched (VSE) medium (Hanisak and Harlin, 1978; Hanisak, 1979; Guiry and Cunningham, 1984). The culture medium was exchanged every 2-3 days and continuously aerated by a filter pump to keep air equilibrium. *Codium sp.* tissue samples were randomly collected for genetic barcoding and species identification.

### 2.2 Experimental incubations under different N sources and concentrations

The experimental setup was carried out with cleaned healthy apex portions. After a 2-day healing period, in total 168 branches were randomly selected and placed into low (5 μM) and enriched (100 μM) N sources (NH□□; NO□□; urea) for another 2 days to obtain *C. cylindricum* thalli with different initial N status. Experimental N concentrations (in its different forms) were selected based on mean environmental conditions experienced by *C. cylindricum* in Hong Kong (20 Years of Marine Water Quality Monitoring in Hong Kong from Environmental Protection Department: https://www.epd.gov.hk/epd/tc_chi/environmentinhk/water/hkwqrc/waterquality/marine-2.html). To achieve the target N status, NH_4_^+^, NO_3_^-^, and urea were added independently to artificial seawater (Instant Ocean with free N) supplemented with VSE medium. Each N source was applied at two concentrations: 5 μM and 100 μM, resulting in six different N treatment combinations. The concentration of each N source was measured (Mulvenna and Savidge 1992; Goeyens et al. 1998; Koroleff 1970; Mulli and Riley, 1995), and listed in Table S1. NaH_2_PO_4_ (10 μM) was also added to avoid phosphorus limitation. After this pre-experimental incubation, samples were randomly assigned to one of the seven temperatures (5–8–12–17–22–26–29°C), with four independent replicates for each treatment. Branches were gradually acclimated from the pre-incubation temperature (17°C) to each target experimental temperature at a rate of 1.5∼2°C per day. After reaching the target temperature, samples were maintained under experimental conditions for 7 days. Cultivation conditions of light and photoperiod were the same as above, and the medium was daily renewed to maintain N concentrations. Aquariums were randomly rearranged during water changes to maintain uniform temperature and illumination. Temperature and light intensity were recorded continuously. The experimental temperatures represent ecologically relevant conditions experienced by *Codium* in Hong Kong (EPD; available from: https://cd.epic.epd.gov.hk/EPICRIVER/marine/). This temperature range was used to assess the thermal performance curve (TPC; nonlinear reaction norm) and variation in its parameters (i.e. the optimal temperature – T_opt; the thermal breadth – T_br; the maximal performance – r_max; and the upper and lower critical – CT_min and CT_max), driven by differences in N sources and concentrations.

### 2.3 Physiological and biochemical parameters

Relative Growth Rate (RGR) was calculated by weighting thalli at the start and end of the experiments: RGR = [(lnW1-lnW0)/(t1-t0)] × 100%, where W0 and W1 were the FW at times 0 and t, respectively, and t was the time interval. Chlorophyll fluorescence parameters (Fv/Fm) were measured after dark-acclimating samples for 20 minutes under each experimental condition (three replicates per treatment) using a diving PAM (Walz, Germany). Photosynthetic pigments content including chlorophyll a, b, and carotenoids were extracted from ∼0.2 g FW samples using 100% ethyl alcohol, following a modified method by Golan et al. (2015). Soluble proteins were extracted from ∼0.2 g FW samples following the method described by Bradford et al. (1976).

Two core physiological indicators (RGR and *Fv*/*Fm*) were fitted with four common nonlinear models (Gaussian, Quadratic, Weibull, and Beta) via the R packages rTPCS (v1.0.0) and nls.multstart (v1.2.0) (Padfield et al. 2021). The best-fitted model with the lowest AICc (corrected Akaike Information Criterion) (Cavanaugh, 1997) value was selected for each curve. Key parameters included the maximum performance (r_max), the optimal temperature (T_opt), the 80% thermal breadth (T_br80%), the critical thermal minimum (CT_min) and critical thermal maximum (CT_max), and 95% bootstrap confidence intervals.

Linear mixed models (LMM) were conducted using the “lme4” package (Bates et al. 2015) to test for differences in other physiological indicators, where temperature (T), N source (NS), and concentration (C) were independent fixed variables (e.g., fixed=RGR ∼ Temperature * N source * Concentration; random=∼|tank). Normality and homogeneity of variance assumptions were verified through visual inspection of QQ plots. Data were transformed either by log10 or by square root to fulfil the requirements for parametric tests. The slopes and intercepts of each temperature fitting model under different N sources and concentrations were extracted and compared based on the average and 95% confidence intervals.

### 2.4 RNA extraction and transcriptome sequencing

Based on the assessed TPCs of the RGR, which can reflect the overall performance of the algae, three temperatures (12, 22, and 26) were selected for further molecular analysis of gene expression (three temperatures × two concentrations × three N sources × four biological replicates = 72 samples). These temperatures are associated with thermal conditions that influence key physiological changes in *C. cylindricum*. Total RNA was extracted using TransZol Up Plus RNA Kit (TransGen, Beijing, China) following the manufacturer’s protocol. RNA quality and integrity were evaluated using an Agilent 2100 Bioanalyzer (Agilent Technologies, USA). That RNA samples with RIN > 7.0, concentration > 50 ng/μl, and clear peak was adopted for deep sequencing. Transcriptome sequencing follows standard procedures on Illumina HiSeq platform, and 200 bp paired-end reads were generated by Novogene Biotechnology Company (Beijing, China).

### 2.5 Transcriptome data processing and differential expression analysis

Adapters and low-quality reads were removed using fastp (Chen et al. 2018). De novo transcriptome assembly and optimization steps were performed using Trinity v. 2.13.2 (Grabherr et al. 2011), DRAP v1.3 (Cabau *et al*. 2017), and Corset v1.09 (Davidson and Oshlack, 2014). Assembly quality was evaluated by N50 (Grabherr et al. 2011), Bowtie2 v2.4.1 (Langmead and Salzberg, 2012), and Busco v4.1.3 (Manni et al. 2021) by mapping back to the sequencing data and the Chlorophyta reference database.

Open reading frames (ORFs) were predicted from assembled contigs using TransDecoder v5.5.0 (Haas et al. 2013). Functional annotation was performed via eggNOG-mapper v2 (Cantalapiedra et al. 2021) and BLASTP search against the UniProt database (E-value cutoff: 1e-5). Transcript abundance was quantified using RSEM (Li and Dewey, 2011). Differential expression analysis was conducted using “DESeq2” (Love et al. 2014). Principal co-ordinates analysis (PCoA) and Permutational multivariate analysis of variance (PERMANOVA) were performed using the “vegan” package to visualize transcriptome expression patterns and test for significant differences among treatments, respectively. Genes with false discovery rate (FDR) < 0.05 and |log2 (fold change) | > 1 were considered significantly differentially expressed genes (DEGs). Gene ontology (GO) enrichment analysis of the significant DEGs was performed using “clusterProfiler” package.

### 2.6 Gene Co-Expression Network Analysis and Fuzzy C-means Clustering

Weighted Gene Co-Expression Network Analysis (WGCNA) was performed using the “WGCNA” package with a merge cut height value 0.25. Module eigengenes (MEs) were calculated to assess correlations between gene co-expression modules and experimental treatments. Modules showing high and significant correlations with experimental variables were retained for downstream functional analysis. Three-way ANOVA was then conducted on ME values to quantify the independent and interactive effects of nitrogen source, nitrogen concentration, and temperature on key gene modules. Hub genes within pivotal modules were identified based on two criteria: gene significance (GS) > 0.2 and module membership (MM) > 0.8. All identified hub genes were subjected to GO enrichment analysis to explore core regulatory pathways.

Fuzzy C-means clustering was performed on all DEGs (from all the comparisons of temperature, nitrogen sources, and concentration) to classify dynamic gene expression patterns under each combined nitrogen treatment (nitrogen source × concentration). Finally, we constructed a regulatory correlation network linking hub genes identified via WGCNA to characteristic gene expression clusters obtained from fuzzy C-means clustering, to elucidate the core molecular regulatory mechanisms underlying the response of *C. cylindricum* to combined thermal stress and nitrogen treatment.

## 3 Results

### 3.1 Thermal performance curves

The best-fitting model for describing thermal performance varied based on the physiological traits and treatments (Table S2). The RGR exhibited the typical bell curve for thermal performance (Figure 1A). In general, the interplay of nitrogen sources and concentrations modulated changes in the thermal plasticity of algal growth, characterized by shifts in both the vertical (r_max) and the horizontal (T_opt, CT_br, and CT_max) dimensions of the TPC (Figure 1B). At high N concentrations, r_max (2.85) of *C. cylindricum* RGR was significantly lower under NH□□ compared to NO_3_^-^ (r_max: 3.53) or urea (r_max: 3.44) treatments (Figure 1B; Table S3). At low N concentrations, no significant differences among N sources were observed. For NH, the maximum RGR at high concentrations was significantly lower (r_max: 2.85) than at low concentration (r_max: 3.79) (Figure 1B; Table S3). However, for NO□□ and urea, there was no significant influence of concentration on r_max (Figure 1B; Table S3). Similar patterns were observed in T_opt (Figure 1B; Table S3). Furthermore, for urea, the differences in various parameters between high and low concentrations were not significant (Figure 1B; Table S3). Both T_br and CT_max were lower under the urea treatment compared to NH□□ or NO□□ treatments, regardless of the concentration (Figure 1B; Table S3).

**Figure 1.**
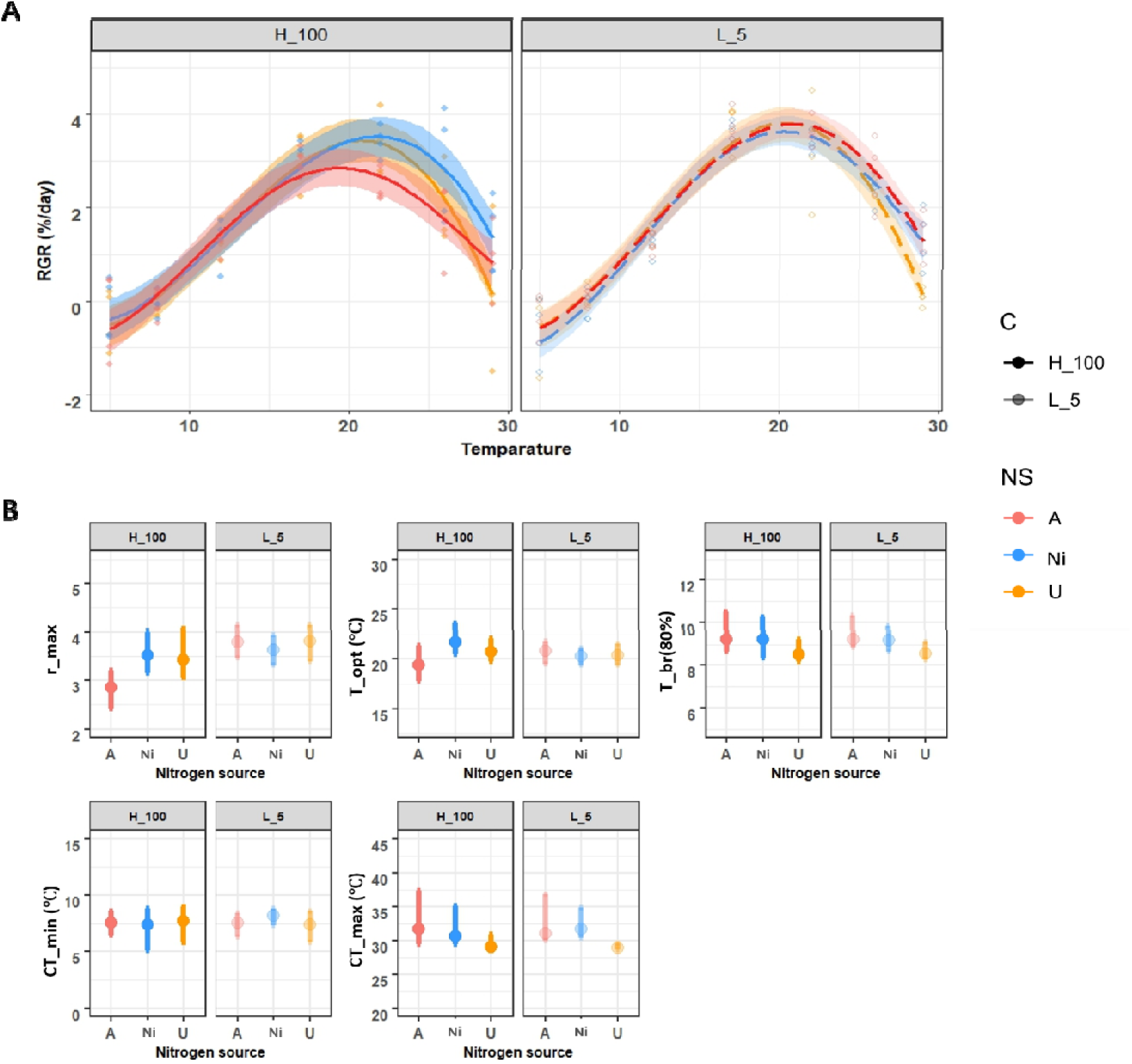
Thermal performance curves (TPCs) of relative growth rate (RGR) for *Codium cylindricum* under different nitrogen treatments. (A) Thermal performance curves and (B) comparison of key TPC parameters. Curves represent RGR performance modeled using the Briere-2 equation (briere2_1999), with shaded ribbons indicating 95% confidence intervals. A, Ni, and U represent NH, NO, and urea, and H_100 and L_5 represent the high (100 μM) and low (5 μM) concentration treatments.

Contrary to the RGR, the maximum quantum yield of photosystem II (Fv/Fm) exhibited linear thermal reaction norms with consistent patterns across all N sources and concentrations, rapidly reaching saturation at temperatures above 10°C (Fig. S1).

### 3.2 Pigments and soluble protein

Photosynthetic pigment concentrations (i.e., chlorophyll a, b, and carotenoids) and soluble protein did not fit any of the non-linear models, showing clear linear relationships with temperature (Figure 2A; Table S4). The slopes and intercepts of these relationships were extracted and compared to assess changes in the influence of temperature driven by N conditions (Figure 2BC; Table S5). Overall, temperature had a positive effect on photosynthetic pigment content (Figure 2A). This relationship was influenced by N sources and concentrations, particularly under high N supply. At high N concentrations, the slopes of temperature–pigment relationships (for Chl a, b, and carotenoids) were significantly higher under NH compared to the NO□□ and urea treatments (Figure 2B; Table S5). However, at low N concentrations, no significant difference in the slopes was detected among N sources. Across all N sources, high concentrations consistently resulted in steeper slopes for the temperature influence on pigment content in comparison with low concentration treatments. The most pronounced effect was observed under NH□□ supply (Figure 2B; Table S5). In contrast, the protein content showed a minor but not significant effect of temperature and its interaction with N sources and concentrations (Figure 2B; Table S5).

**Figure 2.**
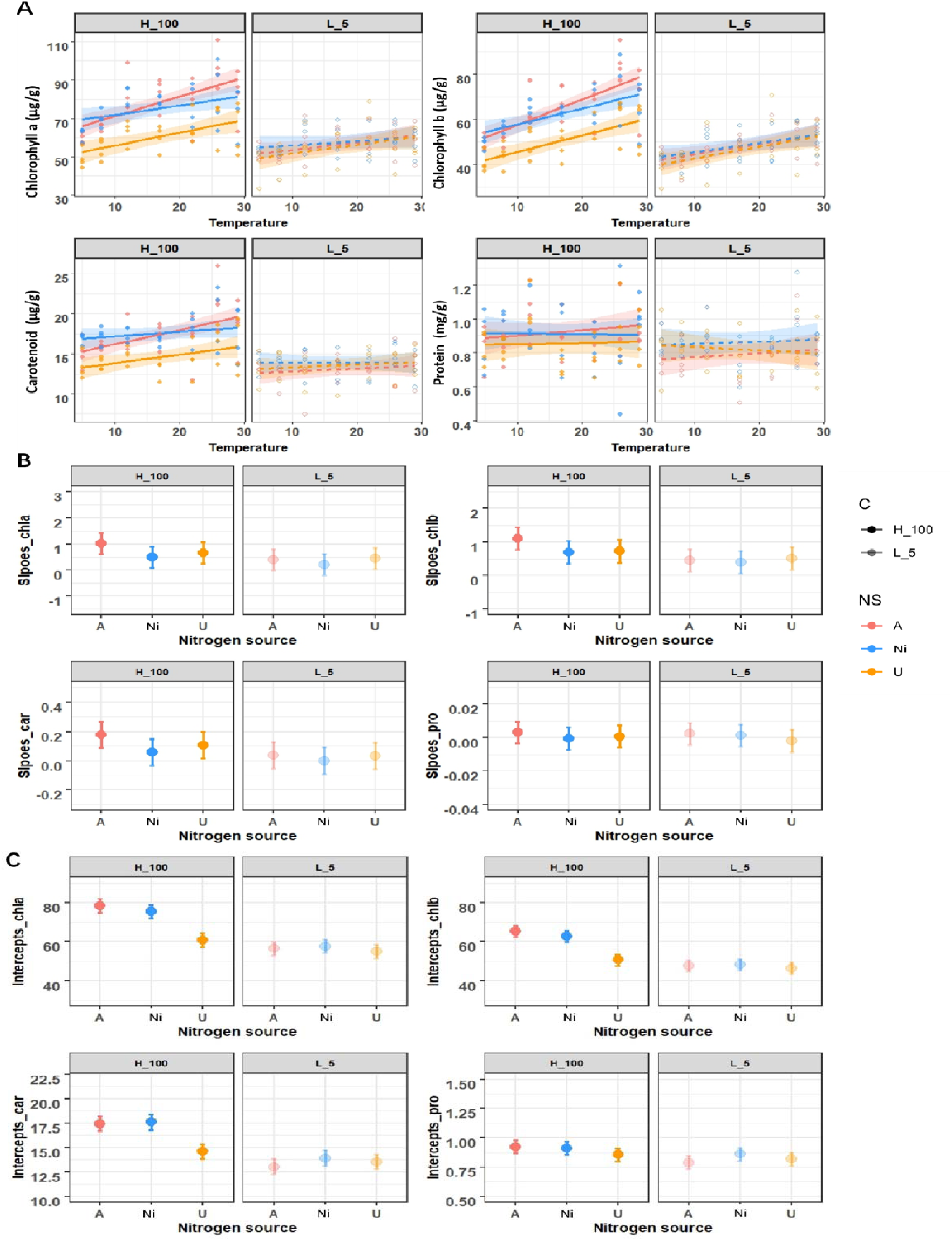
Thermal performance of pigment and soluble protein contents. (A) Linear relationships between temperature and photosynthetic pigments and soluble protein content across different nitrogen (N) sources and concentrations. (B) Comparison of slopes and (C) intercepts derived from the linear regressions. All other symbols, abbreviations, and treatment legends are identical to those described in Figure 1.

At high N concentration, the temperature intercepts of pigments were significantly elevated in NH□□ and NO□□ treatments compared to urea (Figure 2C; Table S5). At low concentration, no significant difference in the temperature intercepts was observed among N sources. As for soluble protein, only NH treatment evidenced significantly higher intercept values at high concentrations than at low concentrations, while no significant differences were observed between concentrations in NO□□ and urea treatments (Figure 2C; Table S5).

### 3.3 Transcriptome analysis and global expression

After quality filtering and assembly, the initial de novo transcriptome assembly comprised 724,305 contigs. Following the removal of chimeric sequences, clustering of contigs, and elimination of redundant or incorrectly assembled transcripts, the final transcriptome assembly consisted of 47,286 transcripts, with an N50 value of 2,476. BUSCO analysis confirmed high completeness, with 93.3% of core Chlorophyta orthologs recovered. On average, 79.2% of RNA-Seq reads mapped back to the assembled transcriptome (Table S6). Various quality metrics indicated that the de novo transcriptome assembly was of sufficiently high quality.

The global expression patterns across treatments were examined using heatmaps and principal coordinate analysis (PCoA), which illustrated the clustering by treatments (Figure S2). Consistent with the physiological data, temperature was the main factor influencing gene expression, with samples from the same temperature grouping together regardless of N source or concentration (Fig. S2). PERMANOVA further confirmed that N source imposed a statistically significant effect on transcriptomic signatures under high nitrogen concentration (*p*=0.019), whereas this differentiation was negligible at low concentration (Figure S2C, Table S7). Based on these global transcriptomic patterns and the physiological results described above, the optimal growth temperature (22°C) was selected for subsequent targeted analyses of nitrogen-driven molecular regulation. This allowed us to minimize the overwhelming effect of temperature and isolate the independent effects of nitrogen source and concentration on algal molecular responses.

### 3.4 N source drives divergent transcriptional responses at high concentration

To dissect the independent regulatory effects of nitrogen source and concentration, we performed pairwise differential gene expression comparisons across the three nitrogen sources (ammonium, A; nitrate, Ni; urea, U) at the optimal temperature of 22 °C, separately for high (100 μM) and low (5 μM) nitrogen regimes. Under high N concentration, pairwise comparisons among nitrogen sources revealed substantial transcriptional divergence. However, under low N concentrations, the number of differentially expressed genes (DEGs) was markedly smaller (Figure 3A,B). No DEGs were shared across the three pairwise comparisons under either concentration regime, indicating that ammonium, nitrate, and urea triggered largely distinct transcriptional reprogramming even under identical temperature and concentration conditions (Figure 3A).

**Figure 3.**
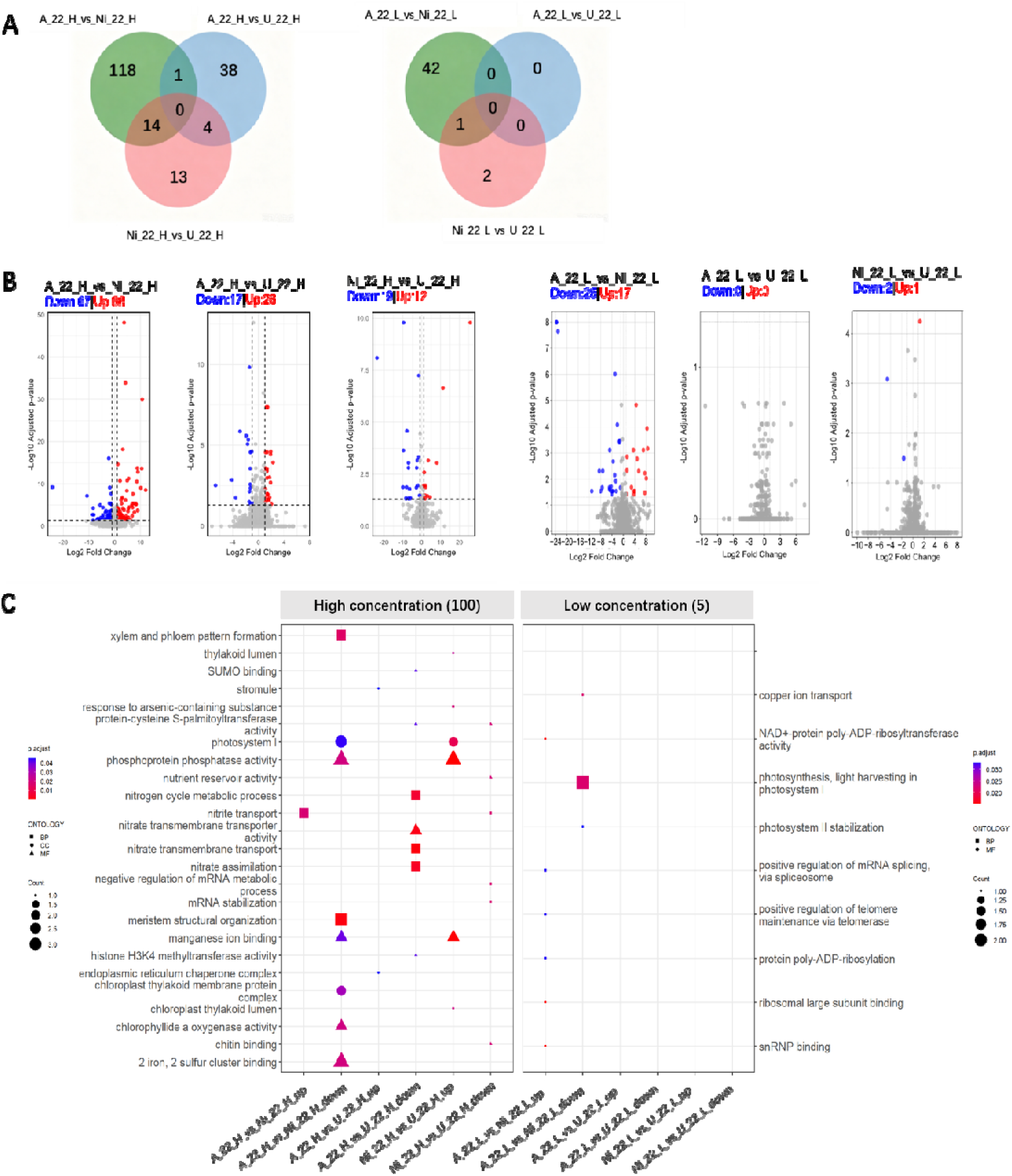
Differentially expressed genes (DEGs) among different nitrogen (N) sources at 22°C. DEGs were identified using thresholds of adjusted p-value < 0.05 < 0.05, |log2FC| > 1). (A) Venn diagram showing the number of shared and unique DEGs. (B) Volcano plot displaying significantly up-and down-regulated DEGs. (C) Gene Ontology (GO) enrichment analysis of DEGs among N sources. BP, Biological Process; MF, Molecular Function; CC, Cellular Component. The *p.adjust* value represents statistical significance controlled for false discovery rate (FDR), and “Count” indicates the number of genes assigned to each specific GO term.

At high nitrogen concentration, pronounced functional divergence was detected among treatments, with ammonium consistently associated with alterations in nitrate and nitrite transport pathways (Figure 3C, Figure S3). In the A vs. Ni comparison, genes encoding formate-nitrite transporters (FNT) were significantly upregulated in the ammonium group. Conversely, multiple pathways were markedly suppressed under ammonium relative to nitrate, including those associated with growth and development (meristem structural organization), photosynthetic machinery (chloroplast thylakoid membrane protein complex, photosystem I, chlorophyllide *a* oxygenase activity, photosynthesis-antenna proteins, Mn ion and 2Fe-2S cluster binding), and phosphoprotein phosphatase activity (Figure 3C; Table S8). In the A vs. U comparison, nitrate-related metabolic pathways (including nitrate transmembrane transport, nitrogen cycle metabolism, and nitrate assimilation) were significantly downregulated in the ammonium group, with the key nitrate transporter gene NRT2.2 coordinately repressed (Figure 3C; Table S8). In the Ni vs. U comparison, each statistically enriched GO term (e.g., nutrient reservoir activity, nitrite transport) contained only one differentially expressed gene, indicating mild functional differentiation between these two nitrogen forms. Nevertheless, photosystem I, phosphoprotein phosphatase activity and manganese ion binding were significantly upregulated in nitrate compared to urea (Figure 3C; Table S8).

In sharp contrast, nitrogen-source-dependent functional differentiation largely disappeared under low N supply. Only weak downregulation of photosystem I light-harvesting functions was detected in ammonium relative to nitrate, and no large-scale pathway reprogramming was observed across nitrogen sources (Figure 3C).

Concentration-dependent effects were most pronounced under ammonium. In this treatment, genes related to nitrate metabolism (e.g., nitrate assimilation, nitrogen cycle metabolic process, nitrate transmembrane transport, and reactive nitrogen species metabolic process) were significantly downregulated at high compared to low concentration, with NRT2.2, NITA, and NRT2.5 among the most prominently affected (Figure 4C; Figure S3B; Table S8). A total of 188, 101, and 5 DEGs were identified for the A_H vs A_L, Ni_H vs Ni_L, and U_H vs U_L comparisons, respectively, with no overlap across the three N-source groups (Figure 4A). In the nitrate group, high concentration was associated with significant upregulation of genes involved in telomere maintenance, chromosome organization, epigenetic regulation, protein phosphatase signaling, and lipid transport, while nitrate transmembrane transporter activity was significantly downregulated (Figure 4C; Table S8). In the urea group, only a small number of DEGs were detected, with significant upregulation of nutrient reservoir activity and lipid transport at high concentration (Figure 4C; Table S8).

**Figure 4.**
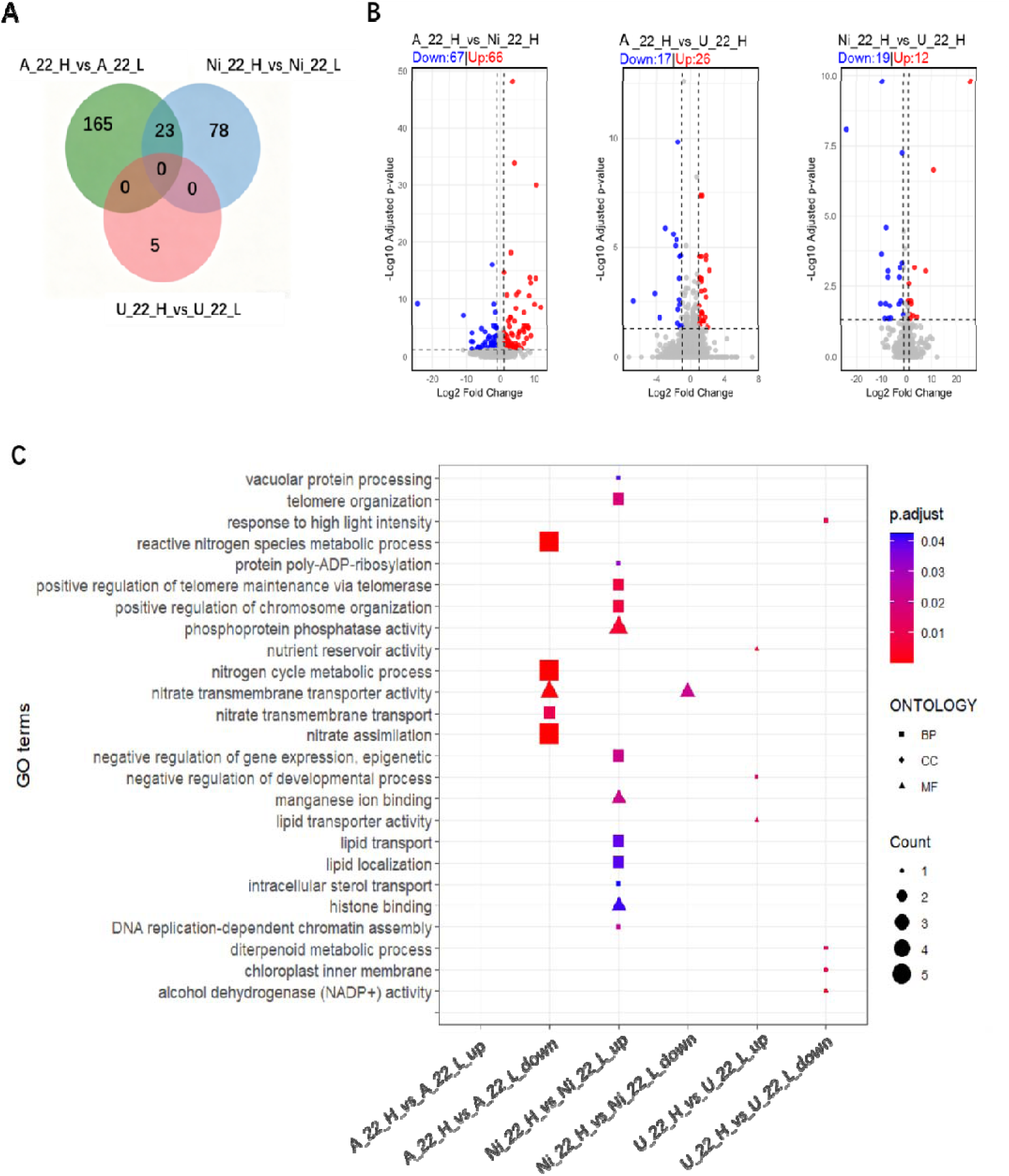
Differentially expressed genes (DEGs) between high and low concentrations at 22°C. (A) Venn diagram showing the number of shared and unique DEGs. (B) Volcano plot showing significantly up-and down-regulated DEGs (adjusted p-value < 0.05, |log2FC| > 1). (C) Gene Ontology (GO) enrichment analysis of DEGs between high and low concentrations. All other symbols and annotations are identical to those in Figure 3.

### 3.5 Nitrogen modulates temperature-driven co-expression modules

Weighted gene co-expression network analysis (WGCNA) was used to identify modules of co-expressed genes associated with experimental treatments (temperatures: 12, 22, and 26°C; N sources, and concentration). A soft threshold power of 18 was selected (R² > 0.80) to construct a scale-free network. After merging modules with a cut height of 0.25, 23 co-expression modules were identified (Figure 5A; Figure S4A). Four modules (MEsalmon, MEdarkorange, MEblue, and MEgreen) showed strong correlations with temperature (|correlation| > 0.5, p < 0.05), underscoring the dominant effect of this factor (Figure 5A). By comparison, only one module was weakly associated with N source (MEsalmon: cor = 0.34, p = 0.004) and one with N concentration (MElightyellow: cor = −0.28, p = 0.02) (Figure 5A).

**Figure 5.**
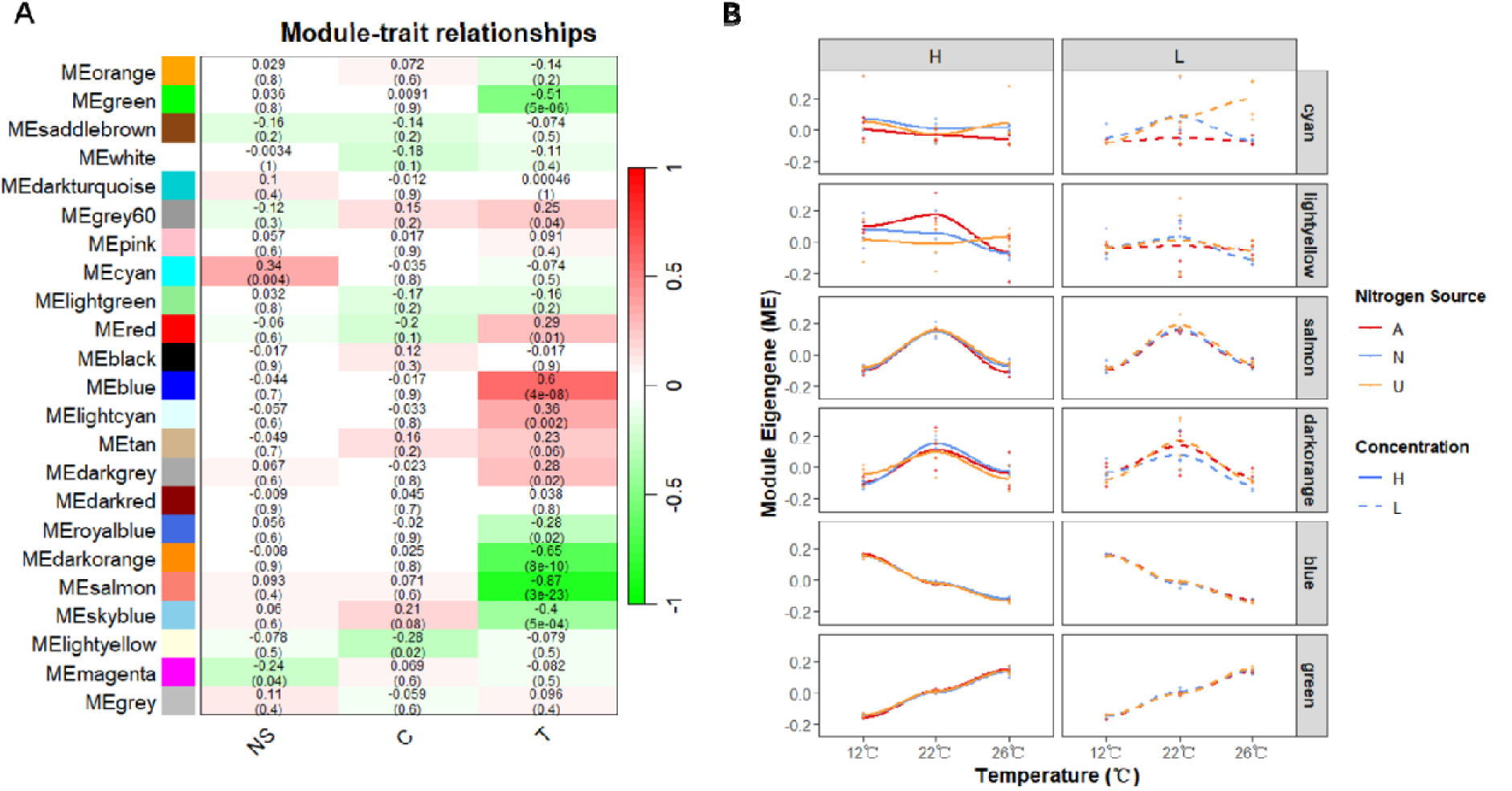
Weighted gene co-expression network analysis (WGCNA). (A) Heatmap displaying Pearson correlation coefficients between module eigengenes and three experimental factors: nitrogen source (NS), nitrogen concentration (C), and temperature (T). Each cell contains the correlation coefficient with its corresponding adjusted p-value in parentheses. (B) Dynamic expression trajectories of six temperature-responsive module eigengenes along the temperature gradient.

We examined module eigengene (ME) and hub gene dynamics across the 12–26°C gradient under different nitrogen sources and concentrations (Figure 5B; Figure S4C). The salmon and green modules exhibited a typical unimodal pattern, peaking at 22°C. The darkorange module appeared modulated by both nitrogen source and concentration. The green and blue modules were continuously up-and downregulated with rising temperature, respectively, and were largely unaffected by nitrogen source or concentration. Three-way ANOVA confirmed significant main effects of temperature on all four temperature-correlated modules (Table S9).

Additionally, both nitrogen source (p = 0.01) and concentration (p = 0.02) significantly affected the salmon module. The cyan and lightyellow modules did not exhibit clear temperature trends but showed differences among nitrogen sources at low and high concentrations, respectively. Notably, a significant interaction between nitrogen source and temperature was detected in the cyan module (Table S9).

GO enrichment analysis was performed on the four temperature-correlated modules to link expression patterns to biological functions (Figure 6A–D; Figure S4B; Table S8). The salmon module was significantly enriched in transcription regulatory activities, including multiple terms related to RNA polymerase II-specific sequence-specific DNA binding, suggesting a coordinated role in transcriptional control and RNA stability. The darkorange module showed highly significant and exclusive enrichment in photosynthesis and light harvesting in photosystem I, indicating a predominant role in photochemical reactions. The blue module was significantly enriched in RNA processing and epigenetic modification pathways, including tRNA metabolic processes, methylation, mRNA splicing via the spliceosome, and histone binding, implicating this module in post-transcriptional regulation and chromatin-related events. The green module displayed significant enrichment in lipid biosynthetic processes (e.g., triglyceride and glycerolipid biosynthesis), chloroplast and plastid membrane components, abscisic acid-mediated signaling, and protein modification and translational regulation (e.g., ubiquitin-protein transferase activity), highlighting its involvement in lipid metabolism, organellar membrane remodeling, and hormonal stress responses under temperature fluctuations. The lightyellow module was enriched in photosystem I-related pathways, while the cyan module showed no significant enrichment (Figure S4B). Together, the salmon module (jointly influenced by temperature and N source, and enriched for transcriptional regulators) and the darkorange module (associated with photosystem I function) represent candidate transcriptional hubs through which nitrogen regime may modulate thermal performance.

**Figure 6.**
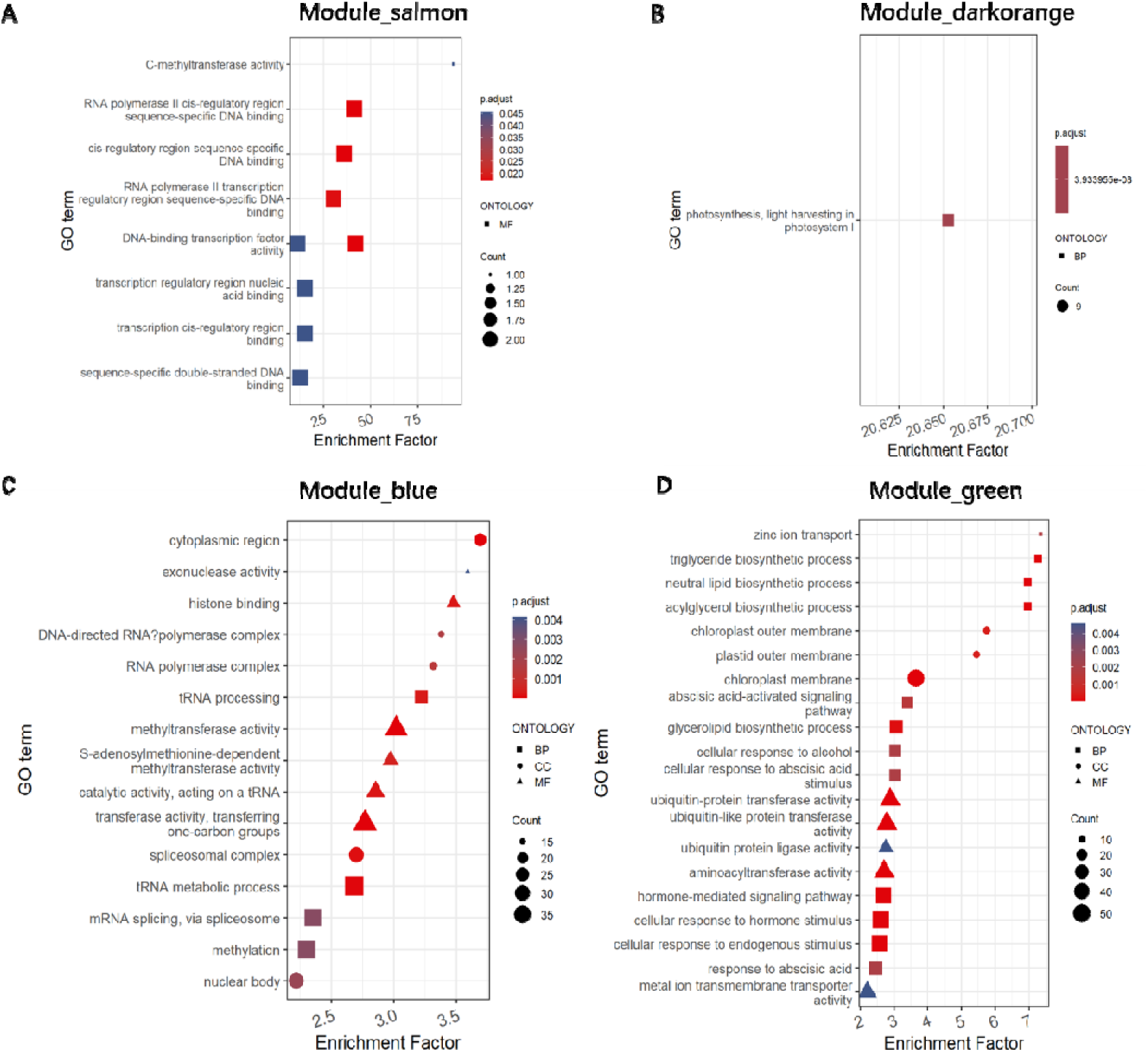
Gene Ontology (GO) functional enrichment analysis of key temperature-associated co-expression modules. (A–D) Top significantly enriched GO terms for four pivotal temperature-correlated modules: (A) salmon, (B) darkorange, (C) blue, and (D) green. All other symbols, abbreviations, and legend definitions are identical to those described in Figure 3.

### 3.6 Nitrogen shifts co-expression module representation across temperatures

Fuzzy C-means clustering was applied to standardized gene expression profiles across the 12–26 °C thermal gradient under all six nitrogen treatments (A_H, A_L, Ni_H, Ni_L, U_H, U_L). Genes with cluster membership values > 0.5 were retained, yielding six distinct transcriptional trajectory types: two temperature-upregulated profiles (up_trend_1, up_trend_2), two temperature-downregulated profiles (down_trend_1, down_trend_2), a V-shaped unimodal trajectory peaking at 22°C (V_trend), and an inverted V-shaped trajectory with maximum expression at 12°C and minimum at 22°C (∧_trend) (Figure 7A,B).

**Figure 7.**
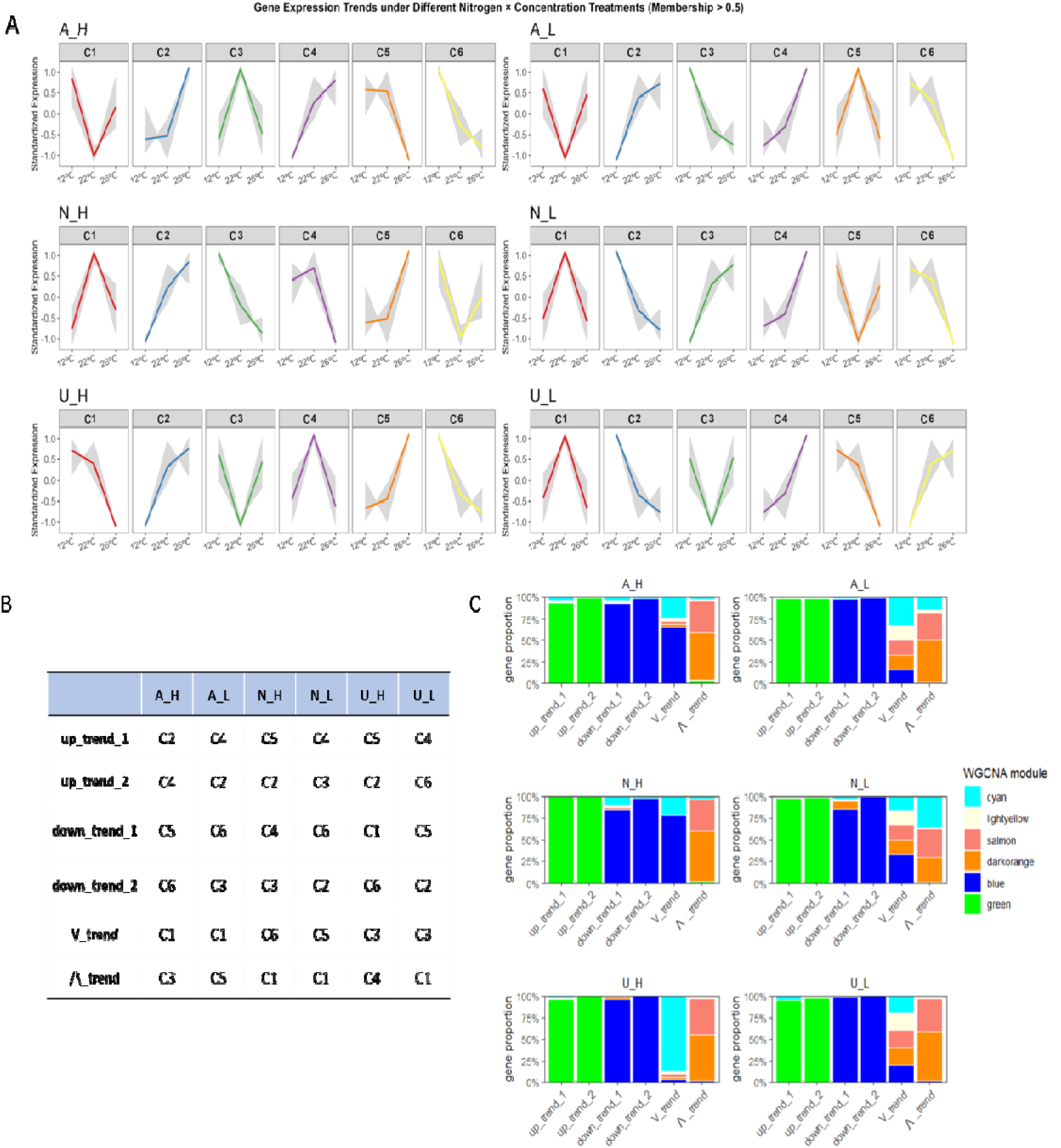
Dynamic trend analysis via Mfuzz clustering and integration with WGCNA modules. (A) Six distinct expression profiles classified by fuzzy c-means clustering across the temperature gradient (12–22–26°C). (B) Classification of the six trend clusters into manual functional categories based on trajectory behavior. (C) Proportion of genes overlapping between static WGCNA module eigengenes and the continuous dynamic gene expression clusters identified by Mfuzz. C1–C6 denote Cluster 1 to Cluster 6, respectively.

Hub genes identified from the WGCNA modules were mapped onto these six expression trajectories, and the relative proportions of hub genes from each module were quantified across the six nitrogen treatments (Figure 7C). The distribution of gene proportions varied among nitrogen treatments (Figure 7; Table S10). Across all treatments, the up-trend gene clusters were predominantly composed of green module genes, whereas down-trend clusters were dominated by blue module genes. The V-shaped unimodal trajectory contained contributions from multiple minor modules. Notably, under low nitrogen conditions, the relative proportions of lightyellow, salmon, and darkorange module genes increased compared to their high-nitrogen counterparts. The inverted V-shaped trajectory (∧_trend) also contained genes from all six modules, with elevated fractions of the cyan module and decreased proportions of salmon and darkorange module genes detected under low nitrogen treatments. These shifts in module representation across nitrogen treatments suggest that nitrogen availability influences the composition of transcriptional programmes deployed along the thermal gradient, particularly modulating the prominence of photosystem-related (darkorange, lightyellow) and transcription-regulatory (salmon) modules.

## 4. Discussion

Environmental temperature plays a critical role in determining the performance, distribution, and fitness of marine macroalgae. However, this influence can be modulated by co-occurring environmental factors, particularly nutrient availability and speciation. Nitrogen (N) is one of the primary macronutrients essential for macroalgal growth and development, constituting key cellular molecules and serving as a pivotal regulator in numerous biological processes (Cai et al. 2012; Shao et al. 2020). Here, we show that N source and concentration jointly determine the thermal threshold at which performance declines, with high ammonium constraining thermal plasticity through a cascade involving the suppression of nitrate assimilation pathways, altered photosystem gene expression, and resource reallocation from growth to pigment accumulation.

### 4.1 N source and concentration shift the thermal threshold for growth

The TPC analysis revealed that both N source and concentration modulated the thermal plasticity of *C. cylindricum*, with effects concentrated in the vertical (r_max) and horizontal (T_opt, T_br, CT_max) dimensions of the curve (Figure 1). At high N concentrations, ammonium significantly reduced maximum growth rate and lowered the thermal optimum compared to nitrate and urea, while nitrate supported the broadest thermal performance. Urea constrained thermal breadth and upper thermal tolerance regardless of concentration. These findings are consistent with evidence from brown macroalgae, where NO enrichment enhances thermal tolerance (Fernández et al. 2020) and buffers thermal stress (Schmid et al. 2020; Colvard and Helmuth, 2017). However, our results extend this understanding to green macroalgae and demonstrate that the effect is not simply a function of N availability; it depends critically on N source.

The observation that high ammonium reduced r_max compared to low ammonium contrasts with the common expectation that increased N availability promotes growth. For Codium species, relatively low nutrient requirements have been documented, with the capacity to store excess N for growth during periods of limited external supply (Chapman, 1998; Hanisak, 1979; Hanisak and Harlin, 1978). High concentrations of NH can inhibit growth in other macroalgae. For example, ammonium concentrations exceeding 2 mg N L ¹ (∼143 µM) significantly inhibited growth of *Sargassum horneri* seedlings (Miki et al. 2016), and even at 60 µM, a concentration lower than that used in our study, long-term NH exposure reduced growth rates in *C. fragile* (Gerard et al. 1990). Our results suggest that the 100 µM NH treatment exceeded the optimal concentration for *C. cylindricum*, triggering stress responses rather than growth enhancement. This represents a concentration-dependent threshold at which ammonium transitions from a nutrient to a stressor. Notably, this threshold appears species-specific: *Ulva australis* shows increased productivity under NH enrichment (Reidenbach et al. 2017), whereas *C. cylindricum* and *S. horneri* (Miki et al. 2016) are inhibited. Understanding the traits governing this threshold across species will be important for predicting macroalgal community responses to coastal nutrient enrichment.

The contrasting thermal responses among N sources also suggest that the widely reported preference of macroalgae for NH, based on lower energetic costs of direct uptake and assimilation (Hurd et al. 2014; Smith et al. 2019; Fan et al. 2020), may not translate into greater thermal resilience. Under thermal variation, nitrate, despite its higher assimilation cost, supported superior performance. This raises the question of whether the energetic saving associated with ammonium uptake is offset by downstream metabolic costs, particularly under thermal stress, when cellular maintenance demands are elevated.

### 4.2 Suppression of nitrate assimilation underlies ammonium-driven thermal constraints

At high concentrations and optimal temperature, ammonium triggered the upregulation of formate-nitrite transporter (FNT) genes relative to nitrate, and the downregulation of nitrate transporter genes (NRT2.1, NRT2.2) relative to urea (Figure 3; Table S8). This pattern suggests that excess NO was exported from cells in response to increased NH availability, consistent with the bidirectional transport capacity of FNT proteins (Lü et al. 2013). The exported unstable NO may be rapidly oxidized to NO and subsequently re-imported via NRT transporters, potentially explaining the upregulation of NRT expression. The downregulation of NRTs in the ammonium-urea comparison further implies the absence of NO under NH treatment and underscores the importance of NO metabolism for *C. cylindricum*.

These transcriptional responses occur against a backdrop of distinct N assimilation pathways. Typically, NH is directly absorbed through ammonium transporters (AMTs) and assimilated into glutamine or glutamate via the GS-GOGAT (Glutamine Synthetase-Glutamate Synthase) cycle (Figure 8) (Inokuchi et al. 2001; Zhang et al. 2022). In contrast, NO must first be reduced to NH through a series of enzymatic reactions involving nitrate transporters (NRT), nitrate reductase (NR), and nitrite reductase (NiR) (Hurd et al. 1995; Young et al. 2005; Islam et al. 2022), while urea must be hydrolysed to NH by urease before assimilation (Smith et al. 2019; Fan et al. 2020; Zhang et al. 2026). Our data suggest that the regulatory control of these pathways, rather than their biochemical efficiency, determines their contribution to thermal performance.

**Figure 8.**
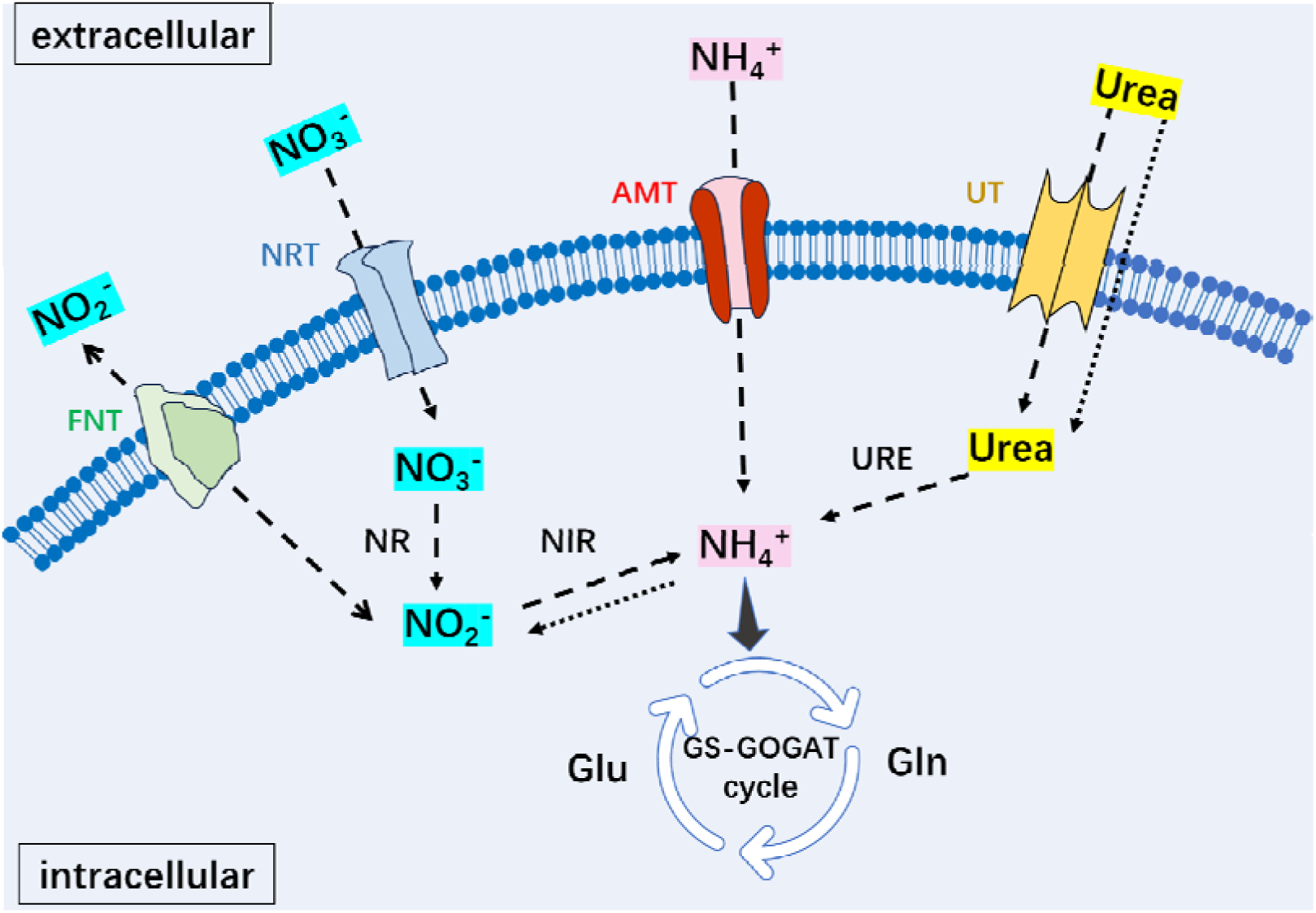
Schematic diagram of the primary metabolic pathways and key enzymes involved in N metabolism in macroalgae. AMT, ammonium transporter; NRT, nitrate transporter; UT, urea transporter; NR, nitrate reductase; NiR, nitrite reductase; URE, urease; GS, glutamine synthetase; GOGAT, glutamate oxoglutarate aminotransferase (glutamate synthase).

Crucially, the suppression of nitrate assimilation genes was concentration-dependent. Comparing high and low ammonium treatments, we observed significant downregulation of NRT2.2, NITA, NRT2.5, and nasB at high concentration (Figure 5; Table S8). This indicates a negative feedback when ammonium is abundant whereby *C. cylindricum* downregulates not only nitrate uptake but also the broader nitrate assimilatory pathway. Previous studies have shown that NH can emit an inhibitory signal that suppresses the expression of genes encoding nitrate transport proteins and related enzymes (Camargo et al. 2007; Galván and Fernández, 2001), and that the presence of NH inhibits NO uptake and utilization in *Ulva prolifera* and *C. fragile* (Hanisak and Harlin, 1978; Head, 1975; Fan et al. 2014). Our results are consistent with this inhibitory model but extend it in an important way. We show that this suppression is not merely a nutritional adjustment, it is associated with reduced thermal tolerance. This suggests that the nitrate assimilation pathway may serve functions beyond N acquisition, potentially contributing to the metabolic flexibility needed to cope with temperature variation.

In contrast, nitrate upregulated photosystem I, phosphoprotein phosphatase activity, and phosphorylation-related genes relative to both ammonium and urea (Figure 3; Table S8). Phosphorylation plays a central role in regulating cellular functions, signaling, and stress metabolism (Damaris et al. 2021), while metal cluster binding proteins (e.g., Fe-S clusters) are essential for cellular respiration, oxygen transport, nucleotide biosynthesis and repair, amino acid biosynthesis, and the synthesis of proteins, cofactors, and vitamins (Dlouhy et al. 2013). The upregulation of these pathways under nitrate suggests that this N source supports the metabolic flexibility needed to maintain performance across a broader thermal range. The concentration comparison further showed that high nitrate upregulated genes involved in chromosome organization and epigenetic regulation (Figure 4-5), indicating additional layers of cellular maintenance that may contribute to thermal tolerance. In contrast, high urea was associated with upregulation of nutrient reservoir activity and lipid transport (Figure 4-5; Table S8), suggesting that when N is supplied as urea, excess nutrients are primarily stored rather than directed toward growth. This storage strategy may explain why urea did not enhance thermal performance despite providing equivalent N: stored nutrients are not immediately available for the metabolic adjustments required under thermal variation.

### 4.3 Resource reallocation from growth to photoprotection under ammonium

While ammonium suppressed growth and shifted the thermal optimum downward, it simultaneously drove greater pigment accumulation with rising temperature. At high concentrations, the temperature–pigment slopes were steeper under ammonium than under nitrate or urea, and ammonium elevated pigment intercepts relative to urea (Figure 2; Table S5). This pattern indicates that under potential NH stress, resources are diverted from growth toward pigment synthesis. In vascular plants, similar trade-offs between growth and photoprotective pigment accumulation have been documented under various abiotic stresses (Demmig Adams and Adams 2006; Li et al. 2018). Our data suggest that an analogous mechanism may operate in *C. cylindricum*, with ammonium stress triggering a reallocation of resources toward photoprotection at the expense of biomass accumulation.

Since pigment synthesis is inherently linked to photosynthetic activity (Martins et al. 2023), we examined the expression of photosynthesis-related genes across N sources. Genes encoding proteins for light harvesting and photosystem function were upregulated under NH and NO conditions (Figure 4d; Table S8), consistent with the increased pigment content observed in these treatments. However, this contrasted with the growth suppression observed under ammonium, suggesting that under NH, the coordination between photosynthetic capacity and growth is disrupted. Rather than translating enhanced light-harvesting potential into biomass, ammonium-treated algae accumulated pigments without a corresponding increase in growth rate. This uncoupling suggests a stress-induced shift in metabolic priorities. Under favourable N sources such as nitrate, enhanced pigment synthesis is coupled to growth, with additional light-harvesting capacity translated into biomass. Under ammonium stress, however, this coupling appears to break down (i.e., pigments accumulate, yet growth remains suppressed). This pattern is consistent with a photoprotective role for pigment accumulation, whereby excess light energy absorbed at elevated temperatures is dissipated before it can damage the photosynthetic apparatus (Figueroa et al. 2009; Lacour et al. 2020).

This divergence between growth and photosynthetic traits highlights the importance of assessing multiple traits when evaluating thermal plasticity. Photosynthesis is a process of light energy conversion, which can rapidly accelerate as temperature increases (Zou et al. 2018). In contrast, growth is a more complex process that, in addition to photosynthesis, involves other physiological activities such as respiration and nutrient absorption, all of which are highly temperature-sensitive and may be limited at higher temperatures (Hurd et al. 2014; Kuebler et al. 1992; Davison et al. 1991). This could explain why optimal growth is typically constrained within a narrow thermal window, above or below which rapid declines in biological reaction rates impact overall performance. Such trait-specific sensitivities to environmental change (Kellermann et al. 2019) underscore why TPCs based on a single trait may not accurately predict overall fitness. Instead, a multi-trait approach is necessary for more accurate predictions of organismal responses to temperature variation. Similar patterns of trait-specific thermal responses have been observed in *Macrocystis pyrifera* (Fernández et al. 2020), and our finding that Fv/Fm showed no N source or concentration effect (Figure S1), in contrast to RGR and pigments, further reinforces this point. Relying solely on fluorescence-based metrics would have masked the N-source-dependent thermal constraints that were clearly evident in growth.

Consistent with this interpretation, the darkorange WGCNA module, enriched for photosystem I and light-harvesting functions, was modulated by both N source and concentration (Figures 5, 6). The salmon module, jointly influenced by temperature and N source, was enriched for transcription regulatory activities (Figures 5, 6), suggesting that N regime may influence thermal responses in part through transcriptional reprogramming rather than purely metabolic adjustment (Sheng et al. 2025). These modules may therefore represent the transcriptional interface through which N regime coordinates the balance between growth-related and photoprotective processes. The fuzzy C-means clustering further revealed that under low nitrogen, the proportions of photosystem-related modules (darkorange, lightyellow) increased relative to high-nitrogen treatments (Figure 7; Table S10). Given that N limitation is known to trigger widespread metabolic restructuring in macroalgae (Hurd et al. 2014), this shift suggests that it redirects transcriptional programmes toward photosynthetic maintenance at the expense of growth, much like ammonium excess. That both N deprivation and ammonium surplus produce similar transcriptional signatures (i.e., increased representation of photosystem modules) suggests a common stress response pathway that warrants further investigation.

### 4.4 N-source-specific regulation of thermal responses

In response to temperature changes, N sources induced shared and distinct transcriptional responses. Core temperature-responsive genes across all N sources were enriched in processes such as ion transmembrane transport, ether lipid metabolism, methyltransferase activity, cell wall morphology, and oxidoreductase activity (Figures 6, 7; Figures S3, S4, S5). Some of these processes, particularly ether lipid metabolism, are closely associated with the fluidity of cellular membranes, which is fundamental for the cellular physiology of marine macroalgae (Schmid et al. 2020). N can modulate membrane fluidity and composition in response to temperature fluctuations, enabling cellular adaptation to temperature stress (Cook et al. 2021; Schmid et al. 2020). The conservation of these responses across all three N sources suggests that they represent a core thermal acclimation programme in *C. cylindricum* that operates independently of N form.

Beyond these shared responses, N sources elicited distinct regulatory programmes that reveal N-source-specific vulnerabilities and capacities. Under low temperature, ammonium specifically upregulated DNA polymerase activity and chlorophyllide-a oxygenase activity, while downregulating processes related to chloroplast membrane composition and mitotic spindle assembly (Figure 6). The chloroplast membrane is a crucial site for photosynthesis, electron transport, and energy synthesis (Song et al. 2021), while the mitotic spindle pole plays a vital role in controlling cell division and the cell cycle (Pavin et al. 2021). These results suggest that ammonium may inhibit growth under low-temperature conditions by disrupting chloroplast membrane function and interfering with cell division. Under high temperature, ammonium-treated algae showed upregulation of nucleic acid metabolism (including deoxyribonucleotide catabolic processes) and downregulation of fat-soluble vitamin and xanthophyll metabolic processes (Figure 7), indicating limited regulatory capacity to withstand elevated temperatures. The downregulation of xanthophyll metabolism is particularly notable, as xanthophylls play key roles in photoprotection and thermal energy dissipation (Goss & Jakob, 2010). Its suppression under ammonium at high temperature may leave the photosynthetic apparatus vulnerable to heat-induced damage.

In contrast, nitrate-treated algae responded to high-temperature stress by suppressing oxidoreductase reactions (Figure 7), reflecting a greater ability to adjust metabolism under thermal challenge. Rather than degrading nucleic acids and downregulating protective pigments, as observed under ammonium, nitrate-treated algae appeared to modulate electron transport and redox balance. These N-source-specific transcriptional programmes provide a molecular basis for the differential thermal plasticity observed at the physiological level, and they suggest that the choice of N form has consequences not only for growth but for how macroalgae cope with temperature extremes.

### 4.5 A threshold model: N source and concentration determine thermal performance limits

Taken together, our results support a model in which N source and concentration jointly determine the thermal threshold at which the performance of *C. cylindricum* declines. When N is supplied as nitrate, active nitrate assimilation pathways support metabolic processes (e.g., photosystem function, phosphorylation, and cellular maintenance) that sustain growth across a broad thermal range. Nitrate also supports membrane-related and enzymatic adjustments that enable more effective responses to both low and high temperature extremes. When N is supplied as ammonium at high concentrations, nitrate assimilation is suppressed, photosystem-related gene expression is altered, and resources are reallocated from growth to pigment accumulation. The consequence is a downward shift in the thermal optimum, reduced maximum growth, and constrained thermal plasticity. Urea limits thermal breadth without strongly affecting the thermal optimum, suggesting a more restricted thermal niche that may reflect the additional metabolic steps required for urea hydrolysis and assimilation.

This model positions nitrate assimilation as a central hub linking N nutrition to thermal physiology in *C. cylindricum*. While our transcriptomic data are correlative and do not establish causality, the consistent association between nitrate pathway expression and thermal performance across N sources and concentrations provides strong circumstantial support for this regulatory cascade. Targeted manipulation of key genes in the nitrate assimilation pathway (e.g., NRT2.2, NITA) would be required to confirm the causal role proposed here.

### 4.6 Ecological implications and future directions

Coastal nitrogen regimes are shifting globally due to eutrophication, aquaculture expansion, and changing land-use patterns (Hutchins & Capone, 2022). These changes alter not only total N loads but also the relative availability of N forms. Ammonium concentrations are rising in many coastal systems, often exceeding nitrate in anthropogenically influenced waters (Glibert et al. 2016). Our findings suggest that such shifts may have underappreciated consequences for macroalgal thermal resilience. If high ammonium availability constrains thermal plasticity and lowers thermal optima, macroalgal populations in ammonium-enriched habitats may be more vulnerable to ocean warming and marine heatwaves than those in nitrate-dominated systems. This vulnerability would be compounded by the fact that ammonium enrichment simultaneously promotes pigment accumulation but not growth, potentially masking stress until thermal extremes expose the underlying physiological constraints.

This has particular relevance for Codium species, which form blooms in nutrient-enriched subtropical and tropical habitats (Lapointe, 1997, 2005; Israel et al. 2010; Chang et al. 2010). While nutrient enrichment may initially promote Codium proliferation, our results suggest that the form of that enrichment, not just its magnitude, will determine how these populations respond to rising temperatures. A shift toward ammonium-dominated nutrient regimes could render Codium populations more thermally sensitive, even as total N availability increases. The relatively low N requirements of Codium (Chapman, 1998; Hanisak, 1979) further imply that moderate nitrate availability may be more beneficial than abundant ammonium for sustaining thermal performance. Future work should test whether the ammonium-driven thermal constraints observed here apply to other Codium species and populations, and whether acclimation or adaptation can shift the concentration threshold at which ammonium becomes inhibitory. Additionally, field-based studies incorporating natural fluctuations in N form and concentration, as well as recovery dynamics following thermal stress, will be essential for translating these laboratory findings to ecological predictions. Understanding how N speciation interacts with temperature to shape macroalgal performance will be critical for predicting the resilience of coastal primary producers under the concurrent pressures of eutrophication and ocean warming.

## Acknowledgements

This research was funded by the grant from the Hong Kong Research Grants Council (17113221) awarded to Juan Diego Gaitan-Espitia.

## Author contributions

Kaile Zhong: conceptualization, methodology, formal analysis and writing—original draft preparation. Juan Diego Gaitan-Espitia: conceptualization, methodology, funding acquisition, supervision, and writing-review & editing. Pamela Fernandez: writing-review & editing. Bayden Russel: supervision, writing-review & editing.

## Ethics Statement

Ethical approval was not required for this study. All samples were collected in accordance with relevant local regulations.

## Data availability

All the transcriptome raw data are available at NCBI: PRJNA1330208 and PRJNA1212827.

